# Suppressive interactions between nearby stimuli in visual cortex reflect crowding

**DOI:** 10.1101/2024.10.07.616799

**Authors:** Leili Soo, Plamen Antonov, Ramakrishna Chakravarthi, Soren Krogh Andersen

## Abstract

Crowding is a phenomenon in which visual object identification is impaired by the close proximity of other stimuli. The neural processes leading to object recognition and its breakdown as seen in crowding are still debated. To assess how crowding affects the processing of stimuli in the visual cortex, we recorded steady-state visual evoked potentials (SSVEPs) elicited by flickering target and flanker stimuli while manipulating the spacing of these stimuli (Experiment 1) as well as target similarity (Experiment 2). Participants performed an orientation discrimination task while accuracy and speed of behavioural responses, along with frequency-tagged SSVEPs elicited by target and flanker stimuli, were recorded. Decreasing target-flanker distance reduced both behavioural performance and target-elicited SSVEP amplitudes. Estimates of the critical spacing, a measure of the spatial extent of crowding, from both behavioural data and SSVEP amplitudes were similar. Additionally, manipulating target similarity affected both measures in the same way. These findings establish a clear connection between the suppression of stimulus processing by nearby flankers in the visual cortex and crowding, and demonstrate the usefulness of SSVEPs in studying the cortical mechanisms of visual crowding.

## Introduction

Visual crowding occurs when objects that are easily recognizable in isolation become difficult to identify in clutter (Andriessen & Bouma, 1976; Bouma, 1970; Strasburger et al., 2011). Since everyday visual scenes are filled with diverse and complex information, crowding affects us at all times and poses a fundamental limit to object recognition (Levi, 2008; Whitney & Levi, 2011). Despite over five decades of extensive research on crowding, its underlying neural mechanisms remain unclear. Gaining a clearer understanding of these mechanisms would allow for the integration of behavioural and neurophysiological perspectives, helping to create a more comprehensive view of how visual information is processed from initial sensory input to final perceptual experience.

The primary manifestation of crowding is spatial interference: the closer the flanking objects are to the target, the worse the ability to recognize the target (Bouma, 1970; Toet & Levi, 1992). This spatial interference is typically measured by critical spacing, the minimum distance at which flankers cease to interfere with target processing (Balas et al., 2009; Bouma, 1970; Pelli et al., 2004). Critical spacing varies depending on various factors (Soo et al., 2018). One such factor that is known to reduce the critical spacing is target-flanker dissimilarity (or ‘pop-out’), whereby a target ungroups from flankers via a featural singleton, such as in shape, polarity, orientation, depth or colour (Andriessen & Bouma, 1976; Chung et al., 2001; Kennedy & Whitaker, 2010; Kooi et al., 1994; Nazir, 1992; Poder & Wagemans, 2007; Scolari et al., 2007). Similarly, critical spacing is reduced when flankers are previewed, i.e. displayed before target onset (Scolari et al., 2007; Watson & Humphreys, 1997), backward-masked (Chakravarthi & Cavanagh, 2009; Wallis & Bex, 2011) or the target location is pre-cued (Albonico et al., 2018; Strasburger & Malania, 2013; Yeshurun & Rashal, 2010). The extent of crowding is larger when flankers have a higher contrast compared to targets (Rashal & Yeshurun, 2014) or if targets are backward-masked (Vickery et al., 2009). Critical spacing can also be decreased with longer display durations or increased with shorter ones (Kooi et al., 1994; Tripathy et al., 2014).

Such behavioural observations have inspired several theories attempting to account for it (Levi, 2008; Manassi & Whitney, 2018; Strasburger et al., 2011; Whitney & Levi, 2011). These include concepts such as spatial pooling, where visual information is integrated over some area (Greenwood et al., 2009; Pelli et al., 2004; Wilkinson et al., 1997); feature mislocalization, where features from distractors are erroneously attributed to the target (Levi & Klein, 1986; Levi et al., 1987; Nandy & Tjan, 2007; Strasburger & Malania, 2013); grouping of nearby visual elements (Manassi et al., 2012); and limitations in attentional resources (Intriligator & Cavanagh, 2001). These theories, while hinting at possible neural processes, cannot pinpoint the specific neural mechanisms responsible for crowding.

Neurophysiological approaches, including functional magnetic resonance imaging (fMRI), electroencephalography (EEG) and magnetoencephalography (MEG), attempt to address the neural mechanisms directly. While the spacing of photoreceptors in the retina are hardwired and impose a physical bottleneck on the richness of detail that can be perceived (Hirsch & Curcio, 1989), the spatial scale of interference seen in crowding is larger than can be accounted for by this sensory bottleneck (Chung & Tjan, 2007; Danilova & Bondarko, 2007; Flom et al., 1963; Tripathy, Srimant & Levi, 1994). Crowding likely occurs in multiple areas along the visual hierarchy (Anderson et al., 2012), from early visual cortex regions like V1 and V2 (Chen et al., 2014; He et al., 2019; Millin et al., 2014) to mid-level areas like V4 (Kim & Pasupathy, 2024) and possibly even higher-order regions in the inferotemporal cortex (Jehee et al., 2007). The exact location and nature of these processes can vary depending on the specific characteristics of the stimuli and the spatial context in which they are presented. There are proposals of fixed interference zones at a cortical level (Pelli & Tillman, 2008; Pelli, 2008), however, this explanation struggles to account for the large variations in critical spacing that are observed when multiple factors that modulate crowding are combined (Soo et al., 2018).

Alternatively, visual crowding can be explained in the context of competitive stimulus interactions. When multiple stimuli are present in the visual field, they compete for neural representation due to the limited processing capacity of the visual system (Desimone & Duncan, 1995; Desimone, 1998). This competition is evidenced by mutual suppression of visually evoked responses, particularly at the level of the receptive field (Beck & Kastner, 2007; Kastner & Ungerleider, 2001). The biased competition model suggests that both bottom-up sensory-driven mechanisms and top-down influences, such as selective attention, can modulate this competition (Beck & Kastner, 2005; Kastner & Ungerleider, 2001). All these aspects share compelling similarities to the crowding phenomenon.

A challenge in studying the neural mechanisms of crowding is that it can be difficult to separate cortical processing of concurrently presented target and flanker stimuli from each other. An elegant solution to this problem is to record “frequency-tagged” steady-state visual evoked potentials (SSVEP). A stimulus flickering at a specific frequency drives a cortical response at the same temporal frequency (Adrian & Matthews, 1934) that can be recorded non-invasively using EEG. When flickering multiple stimuli at different frequencies, each of them will drive an SSVEP response at its specific frequency allowing for a concurrent and continuous measurement of the visual processing of all these stimuli. This method has been utilised in many areas of vision science (reviewed by (Norcia et al., 2015), and has proven particularly powerful in studying visual attention (Andersen et al., 2011b; Morgan et al., 1996). To date, SSVEPs have been employed to investigate crowding only once, showing a crowding-induced suppression of target-elicited SSVEP amplitudes (Chicherov & Herzog, 2015) by nearby flankers. Other studies have investigated suppression of stimulus processing from nearby flanker stimuli using SSVEPs (Fuchs et al., 2008; Keitel et al., 2010, 2013), but none of these studies related these effects to crowding. Thus, the exact relationship between suppressive stimulus interactions and crowding remains unclear.

In two experiments, we investigated how closely the suppression between nearby stimuli reflects the behavioural effects of crowding. We manipulated target-flanker distance (Experiments 1 & 2) and similarity (Experiment 2) while recording frequency-tagged SSVEPs from flickering target and flanker stimuli and behavioural target identification performance. If the recorded SSVEPs capture processes underlying crowding, we would expect corresponding changes in target-elicited SSVEP amplitudes and target identification performance with changes in target-flanker distance and similarity.

## Materials & methods

### Participants

We tested fifteen participants (10 male, 9 right-handed, mean age = 25 years, age range: 21 – 36 years) in Experiment 1 and nineteen participants (15 female, 3 left-handed, mean age = 21.9 years, age range: 18 – 29 years) in Experiment 2. Participants gave written informed consent prior to participation and received £12 or course credits as compensation for their time. All participants reported normal or corrected to normal visual acuity and no known visual or neurological disorders. The experiments were approved by the University of Aberdeen Psychology Ethics Committee (Experiment 1 project number: PEC/3234/2015/6, Experiment 2 project number: PEC/3522/2016/10).

A total of two participants were excluded from statistical analyses because accuracy in the ‘No flanker’ condition was more than three standard deviations below the mean (one in Experiment 1, one in Experiment 2). All participants had at least 50% of all epochs remaining after EEG artifact correction in each condition (71.3 ± 14.1% trials remaining on average in Experiment 1 and 71.4 ± 19.7), and nobody was excluded due to excessive EEG artifacts.

The required sample size was computed using the power calculator provided by Gregory Francis http://www1.psych.purdue.edu/~gfrancis/calculators/calculators.html. Based on previous reports of effect size of target-flanker distance (n = 5; average proportion correct and standard deviation in each flanker distance (in degrees of visual angle, targets at 10° of VA: 1.25°: M = 29.8, SD = 1.10; 2.5°: M = 32.8, SD = 8.32; 3.75°: M = 39.8, SD = 16.35; 5°: M = 58.8, SD = 24.42; 6.25°: M = 70.2, SD = 12.22; No flankers: M = 97.6, SD = 1.67) and similarity (n = 5, average difference between same polarity opposite polarity proportion correct = 3.18, SD = 1.51; Kooi et al., 1994) a sample size of 3 yields a power of 80% for target-flanker distance and a sample size of 5 yields a power of 80% for the similarity manipulation. As our study is only the 2^nd^ to investigate crowding using the SSVEP we had no good prior effect size estimates for the SSVEP effects.

### Task and stimuli

The experiments were presented using MATLAB (The MathWorks, Natick, MA) with the Cogent Graphics toolbox (John Romaya, Laboratory of Neurobiology, Wellcome Department of Imaging Neuroscience) on a 19-inch CRT monitor with a resolution of 800 x 600 pixels and a refresh rate of 120 Hz, viewed from a distance of 60 cm.

*Stimuli and procedure.* Stimuli were displayed for 4000 ms, followed by a 1000 ms inter-trial interval (Figure 1). Participants were instructed to fixate a circle presented in the centre of the screen and focus bilaterally on target stimuli (four superimposed letters ‘T’s in four cardinal orientations, 1.4° x 1.4° visual angle) presented at 9° to the left and right of fixation. In different trials, targets were either presented in isolation or surrounded by four identical flankers (above, below, left and right of target) at one of the tested flanker distances. In Experiment 1, flankers were presented at seven different centre-to-centre distance from targets: 1.7°, 2°, 2.5°, 2.9°, 3.5° 4.3° or 5.1° of visual angle. In Experiment 2, flankers were presented at three different distances: 1.7°, 2.5° and 5°. In any given trial, all eight flankers were presented at the same distance from the targets. To generate SSVEPs, the left target flickered at 10 Hz, the right target at 12 Hz, and all flankers at 15 Hz with synchronous onsets of these stimuli every 500 ms. In Experiment 1, only flanker distance was varied, however, in Experiment 2, targets and flankers were presented either with same or opposite contrast polarity.

**Figure 1.**
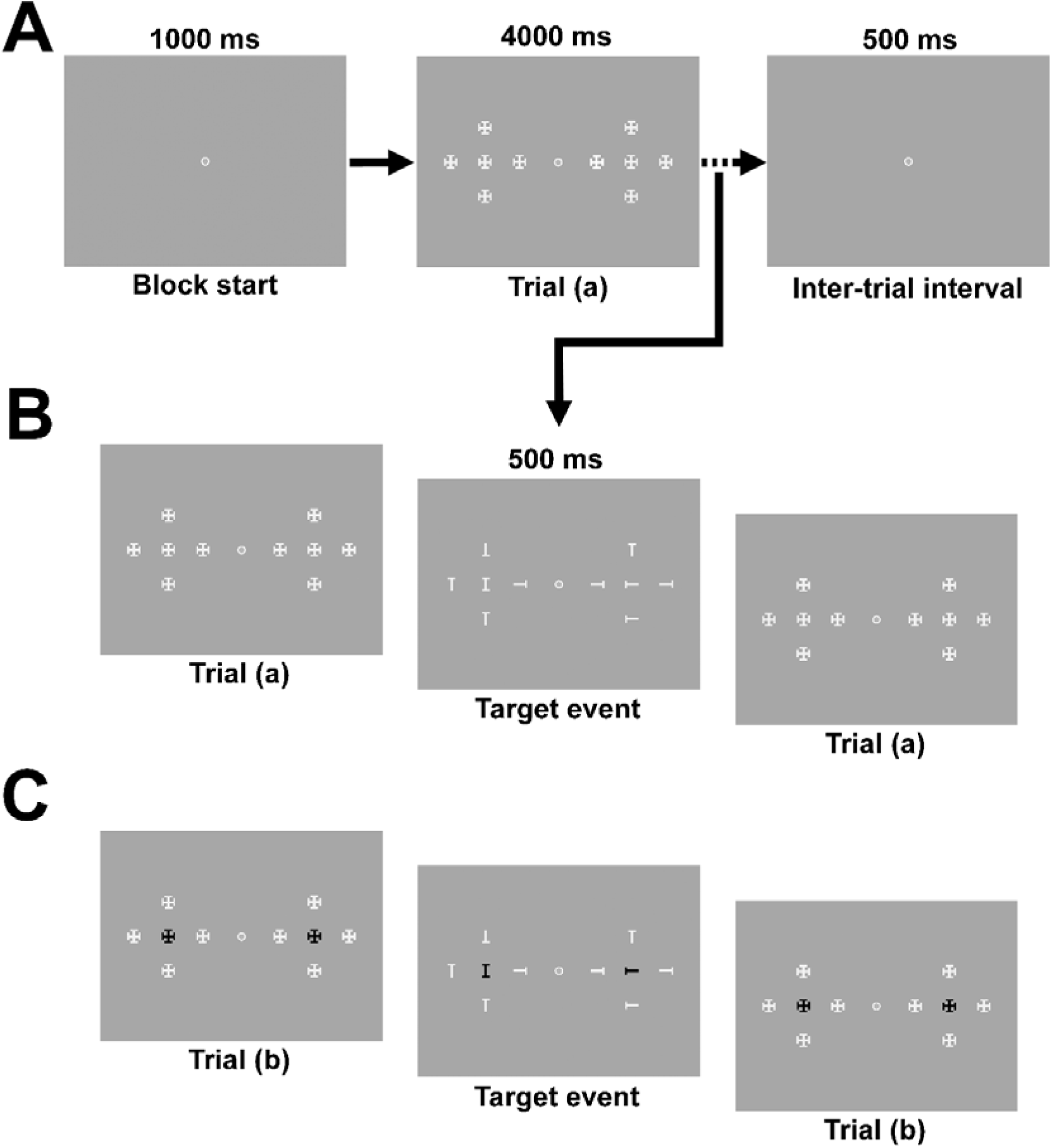
The sequence of events in Experiments 1 and 2. (A) Each block started with a 1000 ms fixation, followed by stimulus presentation for 4000 ms. There was a 200 ms audio-feedback within a 1000 ms inter-trial interval between two consecutive trials. Participants were instructed to attend to bilateral target locations, where shapes were presented either in isolation or surrounded by four flankers on each side at different target-flanker distances. (B) In two thirds of the trials, there were 1-2 target events where a letter ‘I’ was presented in one of the target locations and a letter ‘T’ in the other for 500 ms. Participants were instructed to report the orientation of the target letter ‘T’ (up, down, left, or right) as fast and accurately as possible. Flankers, when presented, were positioned at one of seven (Exp 1) or three (Exp 2) centre-to-centre distances. In Experiment 2, targets and flankers were presented either with the same or opposite contrast polarity (same contrast polarity: (A) and (B), opposite polarity: (C)). In Experiment 2, the fixation circle was presented in colour red isoluminant to the grey background.

*Target events.* During some trials, there were brief 500 ms target events (up to two per trial), where one of the shapes at target locations changed into a target letter T (1.4° x 0.5° visual angle) in one of four orientations (up, down, left, or right) and the other into an I letter (vertical or horizontal; 1.4° x 0.5° visual angle). Simultaneously, all flankers turned into letter T’s presented at random orientations (0°, 90°, 180°, and 270°). Participants were instructed to report the orientation of the letter T at target location by pressing the corresponding arroe key, whilst ignoring the flankers and the letter I. Target event onsets occurred between 1000 to 3500 ms after stimulus display onset, with a minimum of 1500 ms between two target events within a trial. Target events were time-locked to the 500 ms cycles to ensure synchronous onsets of all stimuli.

*Stimulus properties*. In Experiment 1, all stimuli (targets, flankers, and fixation circle) had a luminance of 12.47 cd/m^2^, and the background was 5.61 cd/m^2^. In Experiment 2, the white (positive contrast polarity) stimulus was 11.2 cd/m^2^ and black (negative contrast polarity) stimulus was 0.01 cd/m^2^. At fixation in the centre of the screen was a red circle, isoluminant to the background (5.61 cd/m^2^). The Weber contrast was calculated as:

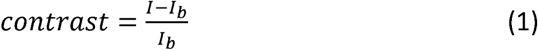

where *I* is the luminance of the stimulus and *I*_b_ is the background luminance, resulting in a contrast of 1.2 in Experiment 1 and ± 1 in Experiment 2.

*Design.* Participants completed between one and three training blocks (20 trials each) to familiarize themselves with the task. In Experiment 1, the main experiment consisted of 576 trials in total (72 trials per condition) divided into 8 blocks of 72 trials. In Experiment 2, the main experiment consisted of 756 trials (54 trials per condition), divided into 6 blocks of 126 trials. Within each block, the conditions were balanced and repeated nine times. Target events occurred in two-thirds of the trials and were randomized to ensure a total of 8 events per condition in each block.

*Feedback and breaks.* At the end of each trial, participants received auditory feedback (high-pitched beep for correct responses, low-pitched beep for incorrect or missed responses). After each block, participants were shown their average accuracy. They were allowed a self-paced break between blocks.

### Behavioural analysis

We analysed the behavioural responses (key presses to indicate the orientation of the target stimulus) that occurred between 300 and 1500 ms after the onset of target events. We classified the responses as correct (participant responded within the time-frame and reported the correct orientation), orientation errors (participant responded within the time-frame with incorrect orientation), misses (there was no response within the time-frame) and false alarms (participant pressed one of the four keys outside the time-frame of preceding target events). The mean proportion of correct responses was calculated for each condition by dividing the number of correct responses by the total number of target events.

### EEG procedure

During the experiment, participants sat in a dark electrically shielded chamber. Participants were instructed to try to blink as little as possible, maintain central fixation and to sit still with relaxed muscles. Electrophysiological data was recorded from 64 scalp electrodes mounted on an elastic cap and 6 additional electrodes (horizontal and vertical EOG and earlobes) using a BioSemi Active-Two amplifier system (BioSemi, Amsterdam, the Netherlands). Scalp electrodes were positioned according to the international 10-20 system (Klem et al., 1999), with electrodes CMS (Common Mode Sense) and DRL (Driven Right Leg) serving as reference and ground during the recording (see https://www.biosemi.com/faq/cms&drl.htm). We used a modified 64-electrode channel layout which omitted electrode positions F5, F6, C7 and C8 in exchange for I1, I2, PO09 and PO10 to prioritize coverage over the occipital lobe. In order to monitor eye movements and blinks, electrooculogram (EOG) was recorded using 4 electrodes, two of which were placed above and below the right eye, and the other two placed laterally to the left and right lateral canthus.

### EEG pre-processing and analysis

EEG data were processed using the EEGLAB toolbox (Delorme & Makeig, 2004) together with custom written routines in MATLAB (The Mathworks, Natick, MA). Epochs were extracted from all trials from 0 to 4000 ms after onset of the flickering stimuli and de-trended (removal of mean and linear drifts). Epochs with eye movements or blinks were rejected from further analysis and all remaining epochs were subsequently subjected to an automated procedure for artifact rejection and correction based on statistical parameters of the data (Junghöfer et al., 2000). On average 71.3 % (Experiment 1) and 71.4 % (Experiment 2) of all epochs remained after artifact correction. All remaining epochs were averaged separately for each of the eight flanker distance conditions and these averages were then subjected to a scalp current density (SCD; λ = 10^-5^, m = 4, 50 iterations) transformation (Pernier et al., 1988; Perrin et al., 1989). SCD transforms differ from scalp potentials in that they are reference free, offer a higher spatial resolution and better correspond with underlying cortical generators (Kayser & Tenke, 2005).

To quantify SSVEP amplitudes, these trial averages were subjected to a Fourier transform. The first 500 ms after the onset of the stimulus display were excluded from this calculation to allow for the SSVEP to build-up. Five electrodes with the highest SSVEP amplitudes averaged across all different target-flanker distances (including the no flanker condition) and participants were selected for each target flicker frequency (10 and 12 Hz) separately. SSVEP amplitudes were averaged across these five electrodes and subsequently rescaled. This rescaling was performed to account for differences in overall SSVEP magnitude between different participants and frequencies (Adamian & Andersen, 2024). Rescaled amplitudes *N_pfc_* were calculated by dividing amplitudes *A_pfc_* by the mean amplitude across conditions (*c*) for each participant (*p*) and each frequency (*f*) separately (Andersen et al., 2011a):

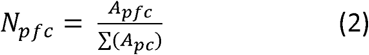

After rescaling, SSVEP amplitudes were collapsed across the two frequencies (i.e., targets).

### Statistical analysis

Behavioural (hits, errors, misses, false alarms, reaction times and proportion correct) and neural measures (target-elicited SSVEP amplitudes) were separately subjected to a one-way repeated-measures ANOVA in Experiment 1 (8 flanker distances including no flanker condition) and a two-way repeated measures ANOVA in Experiment 2 with the factors flanker distance (4 flanker distances including no flanker condition) and target-flanker similarity (no pop-out and pop-out). We used t-tests to compare the behavioural and neural measures between successive flanker distances (Experiments 1 and 2) and flanker contrast conditions using Bonferroni-Holm correction for multiple comparisons.

### Curve Fitting of behavioural and EEG data

In order to estimate critical spacing in Experiment 1, we fitted a cumulative Gaussian function to the mean proportion correct data across trials and target-elicited SSVEP amplitudes as a function of flanker distance (excluding the no flanker condition). SSVEP amplitudes were noisier than the proportion correct data, which precluded fitting curves to individual participant’s SSVEP data. Therefore, curve fits were performed on the group mean performance correct and SSVEP amplitude data. The cumulative Gaussian is given by the following equation (Strasburger, 2001; Wichmann & Hill, 2001):

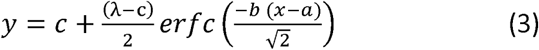

In this equation, y is the proportion correct or target-elicited amplitude, *x* is the flanker distance, *c* is the lower asymptote, λ is the upper asymptote, *b* is the slope of the curve, *a* is the midpoint of the curve, and erfc is the complementary error function. For SSVEP data, the lower asymptote and the upper asymptote were constrained between half of the minimum amplitude across all conditions and to the maximum amplitude across all conditions. For behavioural data, all parameters were constrained to be non-negative (≥0) and the upper asymptote was constrained to a maximum accuracy of 100% (λ = 1). Additionally, for both SSVEP and behavioural data, the midpoint of the curve was constrained between the minimum and maximum flanker distance (1.7 to 5.1); and the slope was constrained between 0 and 10.

The critical spacing x_cs_ is commonly defined as the distance, at which performance reaches 90% of the asymptote:

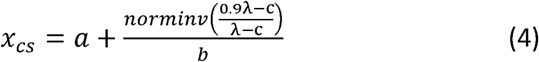

Here, norminv is the inverse of the normal cumulative distribution.

To estimate the sampling distribution of the critical spacings calculated based on proportion correct data and target-elicited SSVEP amplitudes, we used a resampling approach in which 1000 new samples with the original sample size were drawn with replacement from the original dataset. The resulting distributions were then used to calculate the medians and 95% confidence intervals of the critical spacings and of their difference.

## Results

### Behavioural Data

Our two experiments employed a task in which participants identified target stimuli embedded in continuous stimulation. This differs from typical forced choice paradigms commonly employed in psychophysical experiments, such as those on crowding. Whereas responses in forced choice paradigms can be classed as correct or incorrect, it was also possible to commit false alarms and misses in our approach. Our rationale was that if false alarms and misses are (a) rare and (b) largely constant across experimental conditions, then this approach would be sufficiently equivalent to typical forced choice paradigms.

In both experiments, the number of false alarms and misses was indeed fairly low compared to the total number of target events (Figure 2A & 3A). In Experiment 1, the number of misses and false alarms was independent of flanker distance (misses: F(7,104) = 1.11, p = 0.36, η^2^ = 7.9%; false alarms (*F*(7,104) = 2.37, *p* = 0.11, η^2^ = 9.74%). In Experiment 2, there was no effect of either flanker distance (*F*(3,51) = 1.18, *p* = 0.32, *η2* = 4.31%) or similarity (*F*(1,17) = 2.56, *p* = 0.13, *η2* = 1.02%) on misses and there was also no interaction (*F*(3,51) = 1.20, *p* = 0.32, *η2* = 1.68%). There were no effects for the number of false alarms either (flanker distance: *F*(3,51) = 0.74, *p* = 0.53, *η*^2^ = 3.61%; similarity: *F*(1,17) = 0.01, *p* = 0.93, *η*^2^ = 0.00%; flanker distance x similarity: *F*(3,51) = 0.21, *p* = 0.89, *η*^2^ = 0.10%). Accordingly, proportion of correct responses was computed for both experiments and used for all further analyses.

**Figure 2.**
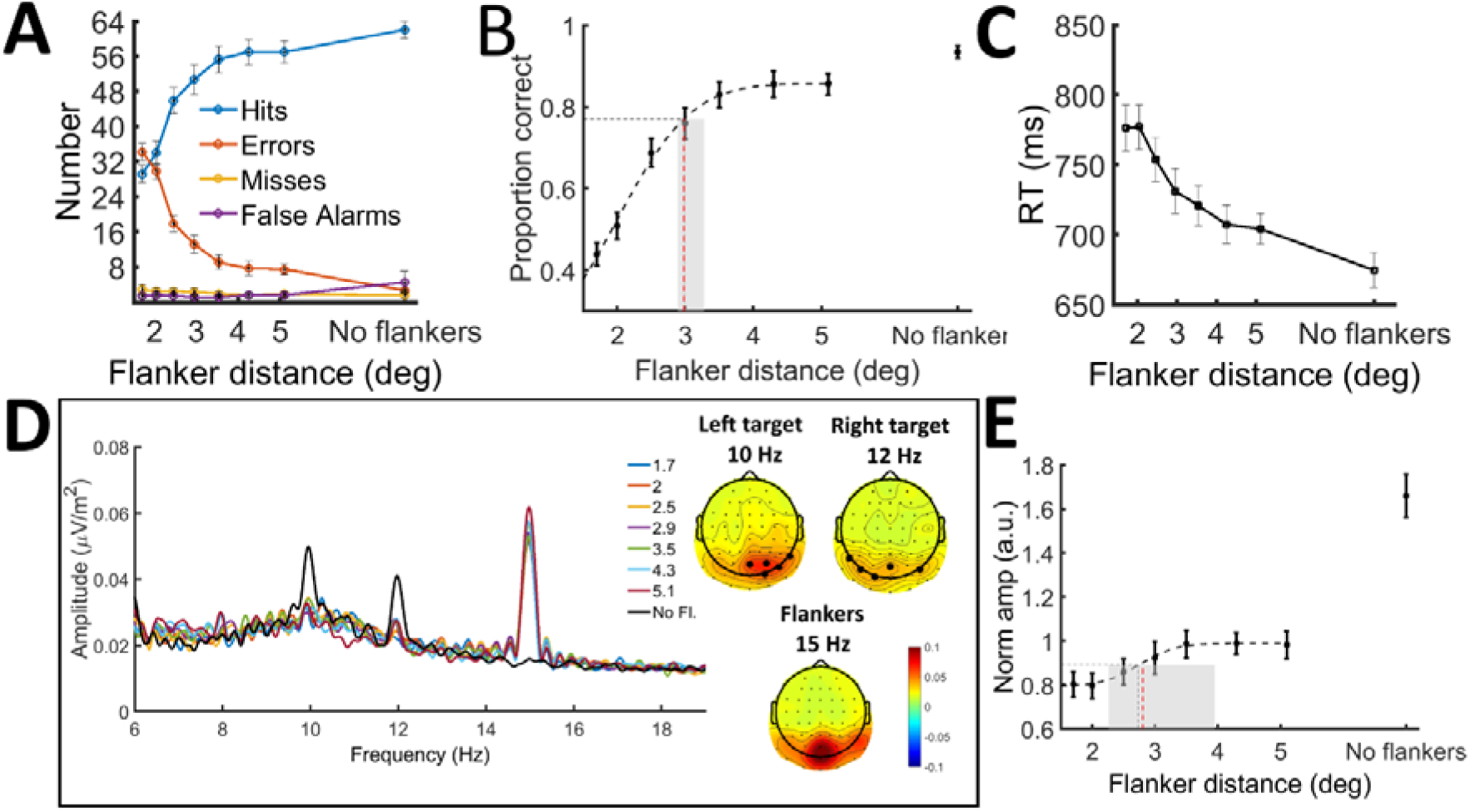
Behavioural and electrophysiological results in Experiment 1. A) The number of hits (correct responses), errors, misses and false alarms as a function of target-flanker distance with standard error of the mean (SEM) from a total of 64 target events per condition. B) Average proportion correct in different target-flanker distances fitted with a Gaussian function. 90% of the asymptote indicated as the critical spacing. Median (red dotted line) and 95% confidence intervals (grey area) of the critical spacings based on 1000 bootstrapped means. C) Average response time in correct trials across participants. D) Grand-average amplitude spectrum obtained by Fourier transform separately for each condition collapsed across occipital electrodes (P04, P08, P6, CP6, P8, P03, P07, P5, CP5, P7 Oz, POz, O2, Iz and O1). Amplitudes peak at simulation frequencies 10, 12 and 15 Hz, however, these peaks are more pronounced when stimulus-specific electrodes are chosen. Topographic maps of SSVEP amplitudes for each stimulus at simulation frequencies averaged across all subjects and conditions (electrode positions chosen for analysis indicated with circles). E) Average normalized target-related amplitudes from five highest amplitude electrodes for each frequency separately, fitted identically to the behavioural data with critical spacing (black dotted line). Median (red dotted line) and 95% confidence intervals (grey area) of the critical spacings based on 1000 bootstrapped means.

**Figure 3.**
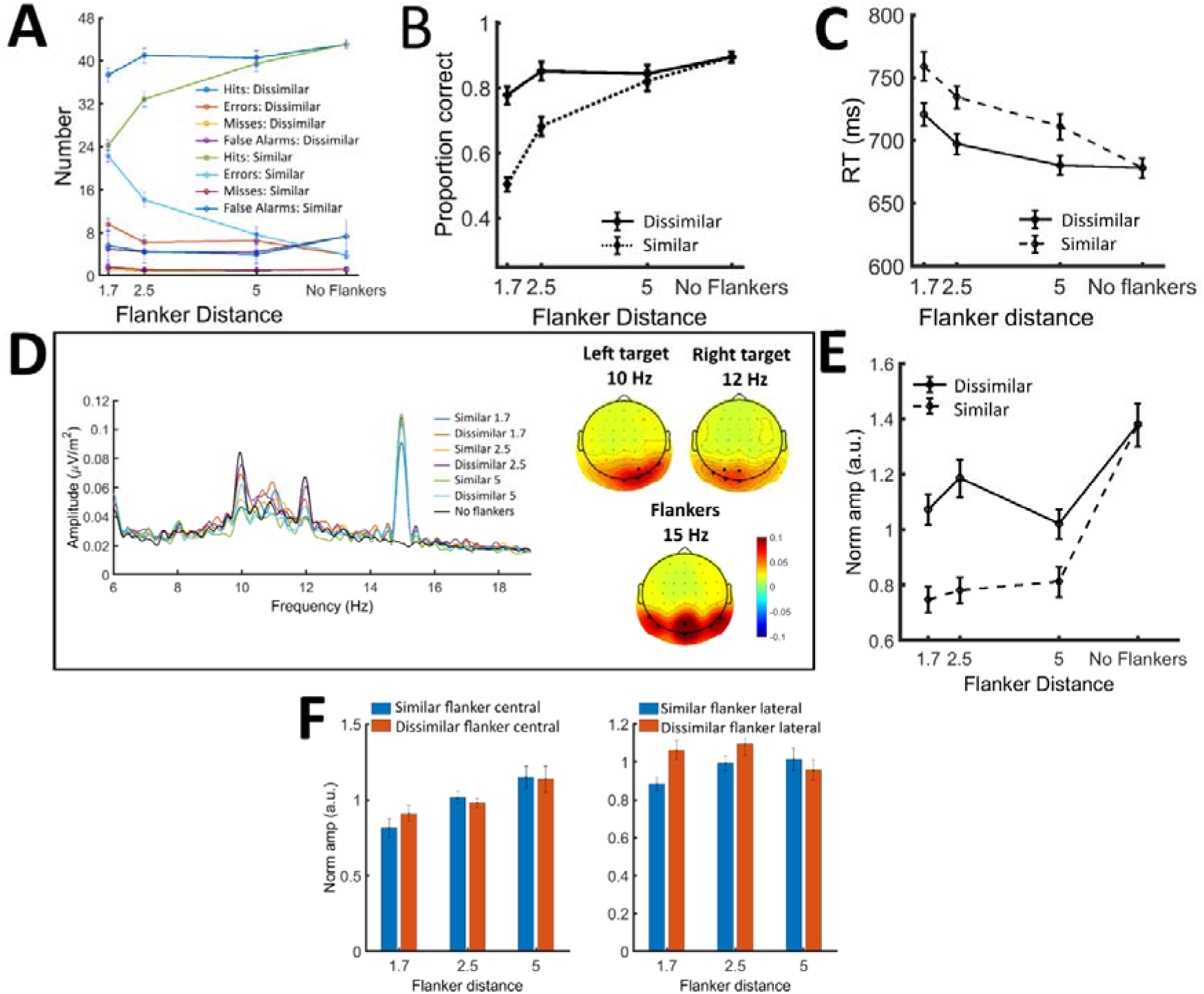
Behavioural and electrophysiological results in Experiment 2. A) The number of hits (correct responses), errors, misses and false alarms as a function of target-flanker distance with standard error of the mean (SEM) from a total of 48 target events per condition. B) The proportion correct as a function of target-flanker distance. C) Average response time in correct trials across participants. D) Grand-average amplitude spectrum obtained by Fourier transform separately for each condition collapsed across occipital electrodes (P04, P08, P6, CP6, P8, P03, P07, P5, CP5, P7 Oz, POz, O2, Iz and O1). Amplitudes peak at simulation frequencies 10, 12 and 15 Hz, however, these peaks are more pronounced when stimulus-specific electrodes are chosen. Topographic maps of SSVEP amplitudes for each stimulus at simulation frequencies averaged across all subjects and conditions (electrode positions chosen for analysis indicated with circles). E) Average normalized target-related amplitudes from five highest amplitude electrodes for each frequency separately. F) Flanker-elicited SSVEP amplitudes separately for central-occipital and lateral-occipital electrode positions in different target-flanker distances with SEM.

In Experiment 1, proportion correct decreased with decreasing flanker distance from the target (*F*(7,104) = 104.22, *p* < 0.001, *η*^2^ = 88.91%; Figure 2B). Pairwise comparisons between neighbouring flanker distances revealed significant decreases between most pairs (1.7°vs. 2°: *t*(13) = 4.07, *p* = 0.001; 2°vs. 2.5°: *t*(13) = 8.85, *p* < 0.001; 2.5°vs. 3°: *t*(13) = 4.22, *p* = 0.001; 3°vs. 3.5°: *t*(13) = 4.37, *p* < 0.001; no flanker vs. 5.1°: *t*(13) = 4.15, *p* = 0.001) as flankers approached the target. Reaction time for correct responses exhibited a marked increase with decreasing target-flanker distance (*F*(7,104) = 28.70, *p* < 0.001, η^2^ = 68.83%; Figure 2C). Pairwise comparisons between neighbouring flanker distances revealed significant increases between no flanker vs. 5.1°: *t*(13) = −4.62, *p* < 0.001.

In Experiment 2, both flanker distance (three distances plus no flanker condition) and similarity (pop-out vs. no pop-out, i.e., opposite vs. same contrast polarity) were manipulated. Proportion correct decreased with decreasing flanker distance (*F*(3,51) = 86.15, *p* < 0.001, *η*^2^ = 48.41; Figure 3B) and this effect was magnified in the same polarity condition (flanker distance x similarity: *F*(3,51) = 64.90, *p* < 0.001, *η*^2^ = 17.04; similarity: (*F*(1,17) = 148.13, *p* < 0.001, *η*^2^ = 18.42). Reaction time for correct responses (Figure 3C) increased with decreasing flanker distance *(F* (3,51) = 68.40, *p* < 0.001, *η2* = 50.40%). Reaction time was lower in the opposite polarity condition (*F*(1,17) = 52.74, *p* < 0.001, *η*^2^ = 16.74%) and there was a significant flanker distance x similarity interaction (*F*(3,51) = 11.09, *p* < 0.001, *η*^2^ = 5.90%) mainly due to the fact that the target-flanker similarity conditions were identical when there were no flankers.

### Target-elicited SSVEP amplitudes

The group average SSVEP amplitude spectrum and topographical maps averaged across all participants and conditions are displayed in Figures 2D and 2E. A one-way ANOVA indicated significant differences in target SSVEP amplitudes across target-flanker distances (*F*(7, 104) = 15.26, *p* < 0.001, *η²* = 53.99%). Notably, there was a very pronounced drop in SSVEP amplitudes between the no flanker and the largest flanker distance conditions (*t*(13) = 5.29, *p* < 0.001, *CI* [0.40, 0.95]).

In Experiment 2, target-elicited SSVEP amplitudes also decreased with decreasing flanker distance (*F*(3, 51) = 13.23, *p* < 0.001, *η²* = 29.26). SSVEP amplitudes were lower when the targets were similar to the flankers compared to when they were dissimilar (*F*(1, 17) = 28.92, *p* < 0.001, *η²* = 10.19%). The reduction in target amplitudes as a function of target-flanker distance was greater in the same contrast polarity condition compared to the opposite contrast polarity condition (interaction: *F*(3, 51) = 6.57, *p* < 0.001, *η²* = 4.73%).

### Critical spacing

Cumulative Gaussian curve fits to proportion correct and target-elicited SSVEP amplitudes averaged across participants produced excellent fits (r^2^ = 0.995 and 0.99, respectively). The resultant estimates of critical spacing were very similar for proportion correct (2.99°) and target-elicited SSVEP amplitudes (2.73°). We employed a resampling approach to obtain estimates of variability of these estimates and their difference. As the pattern in SSVEPs was noisier than in the behavioural data, approximately 5% of the bootstrapped samples did not yield a valid estimate of critical spacing and had to be excluded from our estimates of mean and 95% confidence intervals. The resulting estimates for proportion correct (median = 2.98, 95% CI: [2.89, 3.28]) and SSVEP amplitudes (median = 2.81, 95% CI: [2.26, 3.95]) exhibited substantial overlap, and the distribution of the difference between both included zero (median = 0.14, 95% CI: [-1.05, 0.96]), i.e. it did not provide evidence for a difference.

### Flanker-elicited SSVEP amplitudes

An analysis of flanker-elicited SSVEP amplitudes across flanker distances would not yield unambiguously interpretable results in either experiment, as such comparisons would confound flanker position and distance to the target. Therefore, we exclusively compared flanker-elicited SSVEP amplitudes between the same and opposite polarity conditions in Experiment 2 for each flanker distance separately. Consistent with previous SSVEP experiments in which large symmetric stimuli were presented, the topography of flanker-elicited SSVEP amplitudes exhibited central and lateral peaks (Figure 3D), which most likely reflect activity in early (V1-V3) and later (MT, LOC) visual areas (Andersen, Müller, & Martinovic, 2012). Accordingly, flanker-elicited SSVEP amplitudes were analysed for both clusters separately. At central electrodes, flanker-elicited SSVEP amplitudes did not differ between the same and opposite polarity conditions (1.7°: t(17) = -2.05, p = 0.06; 2.5°: t(17) = 0.84, p = 0.40; 5°: t(17) = 0.41, p = 0.68). At lateral electrodes, flanker-elicited SSVEP amplitudes were larger in the opposite polarity condition than in the same polarity condition at the closest flanker distance (1.7°: t(17) = -3.78, p = 0.001), but there was no effect at the furthest flanker distance (2.5°: t(17) = -1.31, p = 0.21; 5°: t(17) = 1.10, p = 0.28).

## Discussion

The underlying neural mechanisms of object recognition and its breakdown as seen in visual crowding are still unknown. Here, we examined the cortical processing of targets and flankers as a function of the distance (Experiments 1 and 2) and similarity (Experiment 2) between them. The results show that both target identification accuracy as well as target-elicited SSVEP amplitudes are reduced as the distance between targets and flankers reduces (Experiments 1 and 2). This reduction is alleviated when targets are dissimilar to flankers (Experiment 2). Interestingly, the critical spacing estimates for proportion correct and target-elicited SSVEPs yield similar results. This suggests that behavioural measures of crowding and the target-elicited SSVEPs reflect the same phenomenon. The target-elicited SSVEP amplitudes drop as soon as flankers are displayed in furthest away flanker distances (see Figures 3.5B and 3.7D), that is, even when there should be no crowding. This shows that flankers interfere with target processing over large distances in visual cortex. The reduction in flanker-induced processing is alleviated when flankers are opposite polarity to targets. Further, SSVEP amplitudes elicited by flankers are lower at the closest target-flanker distance when they are similar (same contrast polarity) to the target than when they are dissimilar, suggesting that there is mutual interference between similar objects. Overall, these findings suggest that the behavioural breakdown of target identification is reflected in the reduction in target processing in the cortex. The study also points to SSVEPs being a useful tool to studying the cortical processing of multiple objects in the visual field over different spatial scales.

In both experiments, we found that the target-flanker distance did not affect the number of misses or false alarm rates. This shows that target identification but not the detection of target events was impaired, indicating that we measured crowding (Levi et al., 2002; Pelli et al., 2004; Petrov et al., 2007).

The findings can be interpreted in the context of biased competition model of attention (Desimone & Duncan, 1995; Desimone, 1998). The reduction of target processing in the presence flankers, even when the latter are presented at the furthest distance where they are expected to be beyond the critical spacing of crowding, is in line with the proposition that processing resources are spread across the entire visual field for multiple simultaneously presented stimuli (Treue, 2003). Our evidence supports that all objects in the visual field compete for neuronal processing resources (Rolls & Tovee, 1995), possibly due to limitations in available resources (Martin, 1986). This competition for processing resources is influenced by bottom-up mechanisms (here, distance and similarity between targets and flankers). When targets are surrounded by similar flankers, the competition for resources between the stimuli is strong; however, when targets are surrounded by dissimilar flankers, competition is alleviated. These results resonate well with not only the biased competition model of attention but also with the attentional hypothesis of crowding (Fang & He, 2008; He, S. et al., 1996; Intriligator & Cavanagh, 2001; Strasburger et al., 1991; Strasburger, 2005). According to the attentional hypothesis, crowding is caused by coarse spatial resolution of attention (Fang & He, 2008; He, S. et al., 1996; Intriligator & Cavanagh, 2001) or unfocussed spatial attention (Strasburger et al., 1991; Strasburger, 2005). Selection of the target is better when it is dissimilar to the flankers, as it is easier to filter the flankers out despite the coarseness of attentional resolution.

In Experiment 1, we found similar estimates of critical spacing derived from behavioural performance (target orientation accuracy) and those derived from neural measures (target-elicited SSVEPs). As SSVEPs mainly reflect visual processing in V1 (Norcia et al., 2015), our findings suggest that at least part of crowding-related interference occurs in early visual areas. However, target-elicited SSVEPs yielded a larger critical spacing compared to behavioural critical spacing, indicating that crowding might involve interference beyond early visual cortex (Anderson et al., 2012; Freeman et al., 2011). Previous literature suggests that there are two separate hierarchical processes leading to object recognition: object individuation and identification (Intriligator & Cavanagh, 2001; Xu, 2009). Based on this hierarchical separation, critical spacing estimated from the target-elicited SSVEP amplitudes in our data could depict the bottleneck of object individuation, as was proposed previously by Intriligator & Cavanagh (2001). Once objects are individuated during early visual processing, object recognition takes place at a later stage, which might be further affected by interference. This additional interference might be reflected in behavioural measures. This proposal is in line with the evidence that crowding is modulated by object grouping (Chicherov et al., 2014; Ronconi et al., 2016).

While SSVEPs mainly reflect visual processing in early cortical areas, it is not clear whether and by how much SSVEPs are affected by later cortical areas (Norcia et al., 2015). In line with early cortical locus of crowding, previous studies have also shown suppression in V1 (Blake et al., 2006; Chen et al., 2014; Millin et al., 2014). However, the suppression of the target-elicited SSVEP amplitudes does not necessarily reflect bottom-up early suppression of signals. In our paradigm, the targets and flankers were displayed for 4 seconds, within which there were up to two 500 ms target events. The duration of the target display was much longer than commonly used in crowding research; however, this was necessary in our paradigm to allow for the build-up of SSVEPs. Longer display duration has previously been found to decrease the extent of crowding (Kooi et al., 1994; Tripathy et al., 2014) Additionally, in our paradigm, target and flanker locations were previewed by neutral shapes. That is, only the location of flankers was previewed in our paradigm. However, previewing flankers has been found to decrease spatial interference from flankers in crowding only when the actual flanker shapes or identities are previewed (Scolari et al., 2007; Watson & Humphreys, 1997). On the other hand, pre-cuing target locations reduces critical spacing (Albonico et al., 2018; Strasburger & Malania, 2013; Yeshurun & Rashal, 2010). Nevertheless, in our study, both target and flanker locations can be considered to have been pre-cued, which might balance out the target cueing advantage. Hence, while it is not clear whether the suppression in early cortical processing is affected solely by bottom-up or is combined with top-down processing, the suppression is still reflected in the early cortical areas. Our findings, therefore, suggest that target-flanker interference in crowding happens in early visual cortex, which might be augmented by further interference at later stages of processing. This proposal is in agreement with studies proposing that crowding happens in multiple levels of the visual hierarchy (Manassi & Whitney, 2018; Whitney & Levi, 2011).

## Funding Information

This work was supported by the Biotechnology and Biological Sciences Research Council (BBSRC) [grant number BB/J01446X/1].

## Notes

### Competing Interest Statement

The authors have declared no competing interest.

